# Force-dependent facilitated dissociation can generate protein-DNA catch bonds

**DOI:** 10.1101/619171

**Authors:** K. Dahlke, J. Zhao, C.E. Sing, E. J. Banigan

**Affiliations:** University of Illinois at Urbana-Champaign; Massachusetts Institute of Technology

## Abstract

Cellular structures are continually subjected to forces, which may serve as mechanical signals for cells through their effects on biomolecule interaction kinetics. Typically, molecular complexes interact via “slip bonds,” so applied forces accelerate off rates by reducing transition energy barriers. However, biomolecules with multiple dissociation pathways may have considerably more complicated force dependencies. This is the case for DNA-binding proteins that undergo “facilitated dissociation,” in which competitor biomolecules from solution enhance molecular dissociation in a concentration-dependent manner. Using simulations and theory, we develop a generic model that shows that proteins undergoing facilitated dissociation can form an alternative type of molecular bond, known as a “catch bond,” for which applied forces suppress protein dissociation. This occurs because the binding by protein competitors responsible for the facilitated dissociation pathway can be inhibited by applied forces. Within the model, we explore how the force dependence of dissociation is regulated by intrinsic factors, including molecular sensitivity to force and binding geometry, and the extrinsic factor of competitor protein concentration. We find that catch bonds generically emerge when the force dependence of the facilitated unbinding pathway is stronger than that of the spontaneous unbinding pathway. The sharpness of the transition between slip- and catch-bond kinetics depends on the degree to which the protein bends its DNA substrate. These force-dependent kinetics are broadly regulated by the concentration of competitor biomolecules in solution. Thus, the observed catch bond is mechanistically distinct from other known physiological catch bonds because it requires an extrinsic factor – competitor proteins – rather than a specific intrinsic molecular structure. We hypothesize that this mechanism for regulating force-dependent protein dissociation may be used by cells to modulate protein exchange, regulate transcription, and facilitate diffusive search processes.

**Statement of significance:** Mechanotransduction regulates chromatin structure and gene transcription. Forces may be transduced via biomolecular interaction kinetics, particularly, how molecular complexes dissociate under stress. Typically, molecules form “slip bonds” that dissociate more rapidly under tension, but some form “catch bonds” that dissociate more slowly under tension due to their internal structure. We develop a model for a distinct type of catch bond that emerges via an extrinsic factor: protein concentration in solution. Underlying this extrinsic catch bond is “facilitated dissociation,” whereby competing proteins from solution accelerate protein-DNA unbinding by invading the DNA binding site. Forces may suppress invasion, inhibiting dissociation, as for catch bonds. Force-dependent facilitated dissociation can thus govern the kinetics of proteins sensitive to local DNA conformation and mechanical state.

## Introduction

Molecular mechanical forces play an important role in genomic function, governing physiological behaviors including mechanical signaling, gene transcription and expression, and genome structure (1–5). Reaction-rate theory (6–13) and *in vivo* (3, 14–16) and *in vitro* (17–19) experiments suggest that cells may transduce these mechanical stimuli through alterations to ligand-substrate interactions, such as protein-DNA association and dissociation kinetics. For many biomolecular interactions, dissociation requires escaping a single potential energy well, and forces accelerate unbinding by lowering the energy barrier for the dissociation. However, multivalent proteins, *i.e*., those with more than a simple binary set of “on” or “off” states, may dissociate from DNA through multiple, distinct kinetic pathways. These additional kinetic pathways are commonly facilitated by DNA binding by competitor biomolecules from solution (20–23). Because of such “facilitated dissociation” phenomena, the unbinding kinetics of DNA-binding proteins may be considerably more complicated when DNA is subjected to mechanical forces.

Recent work with a variety of biomolecules, such as transcription factors (20, 22, 24–29), metalloregulators (21, 30), chromatin effectors (31), DNA polymerases (32–36), antibody-antigen complexes (37), and other biological complexes (23, 38–48) has demonstrated that facilitated dissociation, in which protein dissociation rates depend on ambient protein or other biomolecule concentration, is a widespread phenomenon both *in vitro* and *in vivo*. Facilitated dissociation contrasts with classical kinetic models of bimolecular complexes that undergo a simple, binary on/off transitions, and have concentration-dependent association rates and concentration-independent dissociation rates. Instead, many proteins undergoing facilitated dissociation are multivalent, so that they may be associated with their substrates in one of several distinct binding states.

As a protein unbinds from DNA via facilitated dissociation, it forms an intermediate state between the fully bound and fully dissociated states, in which it is partially associated with its DNA substrate (Fig. 1A, state P). This partially bound state permits other biomolecules in solution, such as proteins or DNA strands, to compete for contacts with either the protein or the DNA. Subsequent binding by a competitor molecule inhibits the initially bound protein from fully rebinding (Fig. 1A, state S). This accelerates (or “facilitates”) the unbinding of the original protein from its binding site by inhibiting full rebinding of the partially bound protein and providing an additional dissociation pathway (22–24, 28, 42, 49–54). Since spontaneous unbinding without competitor biomolecules may be slow for these protein-DNA complexes (*e.g*., < 10^−3^ s^−1^ for the bacterial protein Fis), facilitated dissociation can lead to as much as a 100-fold enhancement in the dissociation rate at physiological protein concentrations (~ 10 nM for Fis) (20, 22).

**Figure 1:**
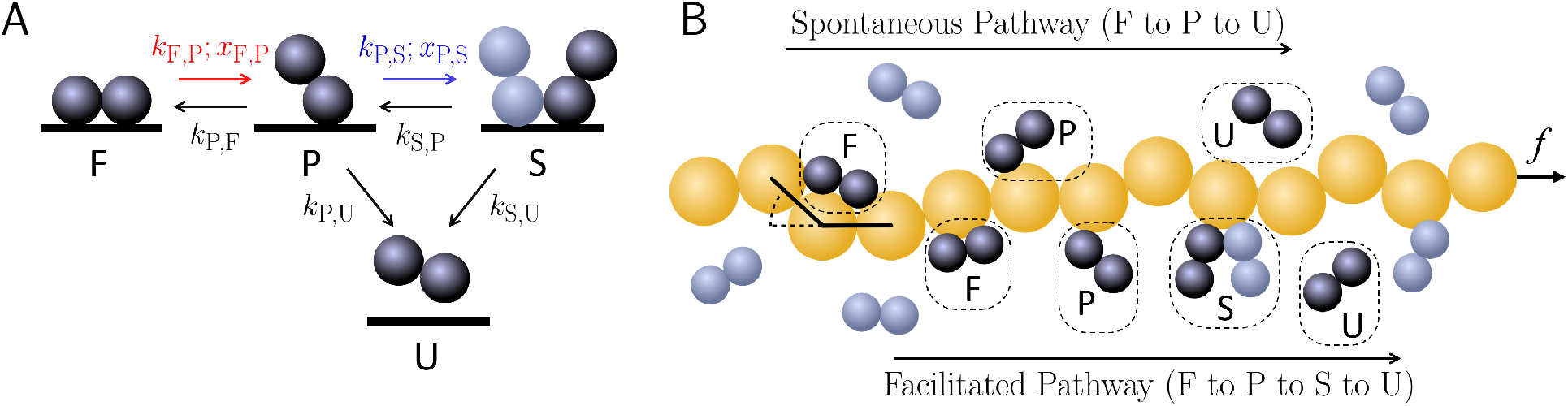
**A**: A schematic of the generic four-state model for protein-DNA dissociation kinetics. The protein-DNA complex may be have single fully bound protein (F), a single partially bound protein (P), two partially bound proteins (saturated, or S), or in the unbound state, no bound protein (U). In the model, the F to P and P to S unbinding steps are force dependent; these steps are indicated with the colored arrows. **B**: A drawing of the many proteins on a long DNA molecule, undergoing the dissociation kinetics depicted in **A**. The top depicts spontaneous dissociation pathway, in which an initially fully bound protein first transitions to the partially bound state, and subsequently, to the unbound state, as shown by the proteins from left to right. The bottom shows the facilitated dissociation pathway, in which an initially fully bound protein transitions to the partially bound state, and then, upon competitor binding, to the saturated state, and finally to the unbound state (from left to right).

Protein-DNA dissociation kinetics may be further modulated by applied forces. *In vivo*, ≳ 1 – 10 pN tension within DNA (or chromatin) can be induced by a number of intracellular and extracellular factors, including transmitted cytoskeletal forces (3, 16), transcription (55–57), DNA replication (58, 59), and chromatin compaction (60–62), among other factors. Since many proteins that undergo facilitated dissociation preferentially associate with bent DNA, flexible DNA, or DNA that is not under tension (28, 63–66), these *in vivo* forces may alter dissociation kinetics through their effects on the geometry of the DNA substrate. Moreover, in these cases, while the total number of proteins bound to DNA decreases under tension *in vitro* (65–67), the kinetics of these tension-dependent proteins are unclear. Many protein-DNA complexes form structures that are disfavored by stretching forces, such as bent or overtwisted DNA conformations; for these complexes, tension should increase the spontaneous dissociation rate of DNA-bound proteins since it lowers the energy barrier for unbinding (6–9, 18). However, since tension suppresses protein binding in this scenario, it should also suppress the facilitated dissociation pathway of protein unbinding. These apparently competing effects suggest that the coupling between force and ligand-substrate geometry may generically lead to unexpected protein-DNA dissociation kinetics within cells.

Indeed, previous work on protein dissociation and unfolding demonstrated that complex force-dependent kinetics can be achieved for structures with multiple transition pathways. For example, for reactions that may occur by two different pathways, inhibition of one pathway can reduce the total reaction rate (13, 68–73). When inhibition is driven by increasing applied force, the bound complex is referred to as a “catch bond” (74, 75). A common conceptual picture for such bonds is that one of the transition states for dissociation resides at a smaller physical distance than the bound state, so that force drives the complex away from that transition (68, 69, 71, 72). However, there are several different generic mechanisms that may underlie such bonds (75). Correspondingly, catch bonds have been experimentally observed for a wide variety of biological proteins and complexes that must withstand physiological forces. These include cellular adhesion proteins in humans (P- and L-selectin and integrin) (69, 70, 76, 77), metazoans (cadherin-catenin-actin) (78), and bacteria (FimH) (79, 80), the molecular motors myosin (81), dynein (82, 83), and PICH (84), microtubule-kinetochore interactions (85), and T-cell receptors under certain conditions (86). In these models and experiments, bond strengthening and slowing of spontaneous dissociation with force are intrinsic properties of the bound complex. In contrast, in the case of facilitated dissociation, it is the *extrinsic* effects of competitor biomolecules that may lead to anomalous kinetics, without bond strength enhancement and suppression of spontaneous dissociation.

We hypothesize that the two dissociation pathways of DNA-binding proteins that undergo facilitated dissociation may together effectively generate catch-bond dissociation kinetics. We predict that when the spontaneous dissociation pathway dominates protein dissociation, increasing force on the DNA substrate enhances the dissociation rate, as is typical of slip bonds. In contrast, when the facilitated dissociation pathway dominates, so that a biomolecule from solution must bind DNA to facilitate unbinding of the DNA-bound protein, the applied force should inhibit dissociation by inhibiting competitor binding; thus, we predict catch-bond kinetics in this regime. We develop a coarse-grained simulation model and a corresponding theoretical model for DNA-bending proteins interacting with DNA, and we explore their force-dependent facilitated dissociation kinetics in the geometry of a common *in vitro* single-molecule experiment (Fig. 1B) (51, 53, 87). Within the model, we identify several distinct types of force-dependent dissociation kinetics: classical slip bonds, catch bonds, delayed onset catch bonds, and force-insensitive bonds. These bonds occur in different physical regimes, depending on the ratio of force sensitivities of each pathway. We investigate how different physical variables, including the preferred binding geometry of the protein-DNA complex and concentration of competitor molecules, may impact force-dependent dissociation. Altogether, the simulations and calculations demonstrate that applied or local mechanical stresses could dramatically modulate the facilitated dissociation of proteins, which could be probed in single-molecule experiments.

## Mathematical and computational models and methods

To explore the consequences of facilitated dissociation in the presence of applied forces, we consider a generic stochastic kinetic model (Fig. 1). There are two essential components of the model: protein unbinding by both spontaneous and facilitated dissociation pathways and inhibition of protein binding by applied forces.

The reaction scheme illustrated in Fig. 1A satisfies the first requirement. In the model, a protein is initially fully bound to DNA (state F), and it must pass from fully to partially bound (state P) before dissociating into the unbound state (state U). While in the partially bound state, a competitor protein may invade the DNA binding site in order to “saturate” it; in this state, the binding site is simultaneously occupied by two partially bound proteins (state S). In the model, competitor binding prevents a partially bound protein from returning to the fully bound state. In experiments, this may also transition the ligand-substrate complex into a highly unstable configuration (22, 54); however, this additional dissociation-enhancing effect is not considered here. In contrast to other schematically similar two-pathway reaction models (68, 69, 71–73), a factor *extrinsic* to the physical state of the bound complex, *i.e*., the concentration of competitor biomolecules in solution, may be tuned to shift the relative weights of the two dissociation pathways. Despite the coarse-grained nature of the model, it has the salient features of facilitated dissociation: two dissociation pathways and extrinsic regulation by competitor proteins. Thus, the reaction scheme describes the unbinding kinetics of various DNA-binding proteins, such as RPA, Fis, and NHP6A (22, 42).

The second condition is achieved by the force dependence of the protein-DNA complex. We specifically consider proteins that bind and bend a long DNA molecule, which is subjected to tension, *f*, as in single-molecule experiments (Fig. 1B) (28, 65, 66, 88–90) or certain in vivo conditions (30). In this setup, when force is applied to DNA, it becomes more difficult to bend, and thus the protein binding affinity should decrease. Therefore, we assume that the rates, *k*_F,P_ and *k*_P,S_, of partial unbinding and competitor binding, respectively, depend on the force applied to the DNA. While we have assumed a specific DNA deformation for the protein-DNA complex, any DNA deformation that is opposed by tension should exhibit qualitatively similar kinetic effects.

We explore this generic model via two complementary approaches, described below. We develop a hybrid Brownian dynamics/kinetic Monte Carlo simulation model to explore force- and concentration-dependent protein dissociation behavior for proteins with different force sensitivities and binding geometries. In parallel, we perform numerical calculations for a theoretical model based on the simulations in order to more easily study the kinetics of individual proteins, analyze the contributions of the two dissociation pathways, and explore regimes of parameter space that are impractical to simulate. Together, the two approaches elucidate how the kinetics described by Fig. 1A may be manifested in the typical experimental scenarios depicted in Fig. 1B.

### Hybrid Brownian dynamics/kinetic Monte Carlo simulations

#### Model for DNA polymer dynamics

We adapt a coarse-grained Brownian dynamics simulation model for DNA-binding proteins binding that demonstrates both spontaneous and facilitated dissociation (Fig. 1B) (53, 87). DNA is represented by a strand of *N* = 100 beads, indexed by *i*, of radius *a* = 4 nm at positions **r**_*i*_. The strand is tethered to a stationary boundary at one end, and a pulling force *f* is applied to the other end. Units are non-dimensionalized as follows: distance with respect to bead radius 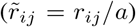, time with respect to the diffusion time of a single bead 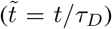, and energy with respect to 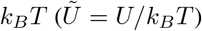. Throughout the text, variables with tildes denote non-dimensionalized quantities.

Neighboring beads in the DNA polymer are connected by Hookean springs, which is governed by the stretching potential:

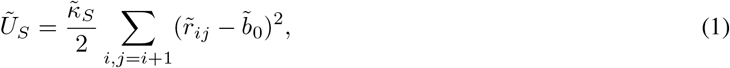

where 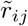 is the distance between adjacent beads *i* and *j*, and 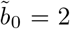 is the equilibrium distance between two beads. 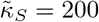 to prevent large deviations of 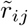 from 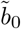.

Excluded volume is included via a shifted Lennard-Jones potential:

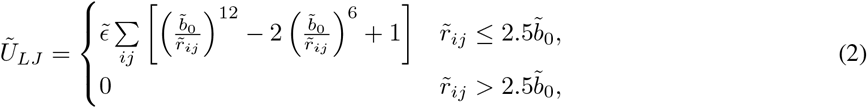

where the magnitude of the potential is controlled by 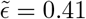, and 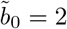 is location of the minimum.

A bending potential is included to maintain the stiffness of DNA as well as to incorporate the effects of the local bending deformations that may be induced by DNA-binding proteins:

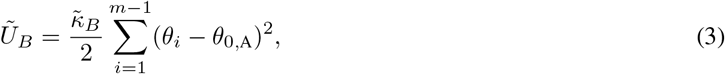

where 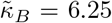 ensures a DNA persistence length of 50 nm, and *θ_i_* is the angle between the bonds of beads *i* and *i* – 1 and beads *i* and *i* + 1. *θ*_0,A_ is the equilibrium angle between these two vectors, and its value depends on whether protein is bound to DNA bead *i*, which is denoted by the state index A.

Monomer dynamics are overdamped and governed by a Langevin equation that includes the three potentials described above:

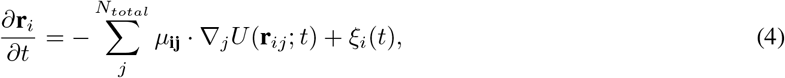

where *μ*_**ij**_ = *δ_ij_**δ***/(6*πηa*) is the freely draining Stokes mobility matrix, *η* is the solvent viscosity, ***δ*** is the identity matrix, *δ_ij_* is the Kronecker delta, *U* = *U_S_* + *U_LJ_* + *U_B_* is the sum of all bead potentials, and *ξ_i_* is a random velocity that satisfies the fluctuation-dissipation theorem, 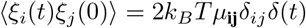.

#### Model for protein binding and unbinding kinetics

Proteins in the system are implicitly modeled as a field of proteins that interacts with the DNA strand. A four-state model is used to describe the possible states of a DNA bead: fully bound (F), partially bound (P), saturated (S), and unbound (U), as shown in Fig. 1A. The partially bound and fully bound states correspond to a single protein interacting with a single DNA bead, while the saturated state corresponds to each of two proteins partially binding the same single DNA bead. DNA in either the unbound (U) or partially bound (P) states locally has an equilibrium bending angle of *θ*_0,U_ = *θ*_0,P_ = 0, while a non-zero equilibrium angle is associated with the fully bound (F) and saturated states (S): *θ*_0,F_ = *θ*_0,S_ > 0. These angles can be changed independently from each other, but are kept the same in this work for simplicity; unless noted, simulation results for *θ*_0,F_ = *θ*_0,S_ = *π*/6 are shown.

Transitions between two states, A and B, are dictated by state-dependent energy barriers, 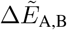. The binding state of a DNA bead, denoted by Ω_*i*_, is updated with a Monte Carlo step every 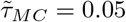. This time step is chosen to ensure that if an (implicit) protein is found to have diffused so that it is near the DNA binding site, the subsequent binding time, *τ_b_*, matches the diffusion time, *τ_D_*, of a single bead (53). A random number, 0 ≤ *ζ* < 1, is generated, and for DNA site *i* in state Ω_*i*_ = A, the binding update occurs as follows:

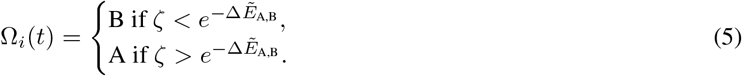

Most transitions in this model have fixed energy barriers for binding or unbinding: 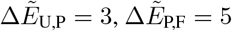 (where the additional 2*k_B_T* accounts for entropy loss upon full binding (87)), and 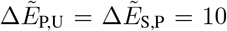. Due to the universal nature of facilitated dissociation, the results are broadly applicable beyond this particular set of energy barrier choices (53).

Two key transitions have energy barriers that vary as a function of force. The form of the energy barrier for the competitor protein binding event to DNA site *i* in the facilitated dissociation pathway (P to S) is:

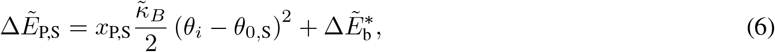

where the second term is a constant energy barrier. The first term in Eq. 6 imposes an energetic penalty for binding that arises from DNA being bent away from the equilibrium bending angle, *θ*_0,S_, of the saturated protein-DNA complex. This indirectly leads to a force dependence for the energy barrier. This indirect force dependence is used because because the force applied to the DNA strand does not directly act on the unbound competitor protein (87). As more force is applied to the DNA strand, the average angle, *θ_i_*, between DNA bead bond vectors increases. Therefore, on average, the DNA binding site is further from its preferred binding geometry at higher applied forces, which decreases the likelihood of protein binding. The force sensitivity parameter, *x*_P,S_ > 0, controls the strength of the effect of force on the binding energy barrier. To ensure that the zero-force total protein dissociation rate, *k*_off_, is the same for all parameters, we include the adjustable parameter 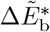 to account for the effects of the force-dependent term on the zero-force off rate. The calibration procedure for 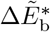 is described in detail in the “Variables and observables” section, below, and the values of 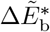 are listed in Table S1.

The energy barrier for the force-dependent partial unbinding step (F to P) has the form:

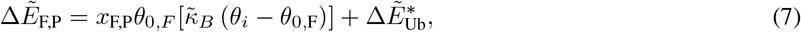

where the second term is a constant unbinding energy barrier. We directly incorporate the effect of force on the protein-DNA complex, which leads to a different functional form of the force-dependent term. The first term is proportional to the derivative of the bending energy, which is quadratic as in Eq. 6 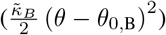. Here, we use a force-like term because the fully bound protein-DNA complex is directly destabilized by tensile forces applied to the DNA strand. Because we must account for local bending conformation of DNA to determine the unbinding energy, the form of the energy barrier in Eq. 7 differs from that of Eq. 6 (87). The coefficients of the first term of Eq. 7 are the force sensitivity parameter, *x*_F,P_, and the equilibrium bending angle, *θ*_0,F_, of the fully bound state. The force sensitivity *x*_F,P_ > 0 controls the strength of the force-dependent term. As in Eq. 6, the last term, 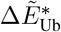, is an adjustable parameter that is calibrated to match 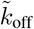 at *f* = 0 for all parameters. The calibration procedure is described below, and the values of 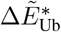 used in this study are listed in Table S2. Physically, we may interpret *x*_F,P_ as a measure of the distance a protein must be displaced in order to partially dissociate, or the position of the transition energy barrier (8–10, 72, 75).

The force dependence of *k*_F,P_ but not *k*_P,U_ models insensitivity of the spontaneous dissociation step to applied forces, while partial dissociation events depend on force. These kinetics are suggested by observations of DNA-binding proteins such as NHP6A, HMGB, and HU (88, 90). Furthermore, the force dependence of the saturation transition models inhibition of protein binding by tension (*e.g*., as in (66)), which we have assumed is relevant to competitor protein binding.

#### Variables and observables

We define the sensitivity ratio as *x*_P,S_/*x*_F,P_, which is a simple measure of the relative sensitivities of each dissociation pathway to force. A high ratio (> 25) indicates that the facilitated pathway is much more sensitive to force, whereas a low ratio (< 10) indicates that the spontaneous pathway is much more sensitive to force. Simulation results for different sensitivity ratios are obtained by fixing *x*_P,S_ = 10 and varying *x*_F,P_.

We measure the off rate, 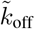, in simulations by counting the number, *n_B_*(*t*), of initially bound proteins that remain bound to the DNA over time. We initialize the DNA strand with every nonspecific binding site in the fully bound state, and then allow these originally bound proteins to unbind and competitors to bind over time. We simulate until all originally bound proteins unbind. We take the average of 100 independent trajectories to obtain 〈*n_B_*(*t*)〉, and thus the mean dissociation rate, 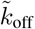.

The energy barriers 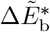 and 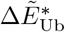 in Eqs. 6 and 7 are calibrated to compensate for the effects of the force sensitivities, *x*_P,S_ and *x*_F,P_, on the measured off rates, 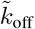. Calibration is necessary because in zero-force simulations, different force sensitivities can lead to different off rates due to differences in how DNA conformational fluctuations are coupled to protein binding and unbinding. In experiments, these differences would be interpreted as the result of different energy barriers. Thus, changes in force sensitivities in experiments would only be detectable as changes in the force dependence of 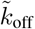. By calibrating 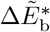 and 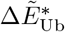, the same is true of the simulations.

To calibrate the energy barrier, 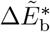, for competitor binding, we fix *x*_F,P_ = 0 and 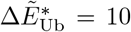 and measure 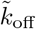 for different values of *x*_P,S_ and 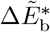 at fixed concentration (*c* = 100 *μ*M) and force (*f* = 0 pN). For each *x*_P,S_, we identify the value of 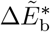 for which 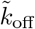 equals a reference value (taken as 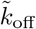 for *x*_P,S_ = 10 and 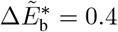).

Then, to calibrate 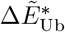, we fix *x*_P,S_ = 10 and 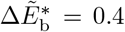, and for each of *x*_F,P_, we measure 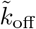 for different values of 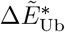 at fixed concentration (*c* = 100 *μ*M) and force (*f* = 0 pN). For each *x*_F,P_, we identify the value of 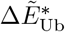 for which 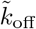 is equal to the reference off rate.

Classification of force-dependent kinetics is done by determining whether *k*_off_ increases or decreases with increasing force, *f*. Increasing 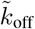 indicates a slip bond, while decreasing 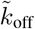 indicates a catch bond. In some regimes, 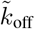 is insensitive to force. We identify such regimes by 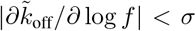. This cutoff, *σ* = 1.5· 10^−6^, is empirically chosen to identify regions where 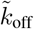 varies slowly with *f*, but is not a transition between slip and catch bonds; the cutoff also removes artifacts from regimes in which statistical noise overshadows small differences in 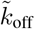.

### Theoretical geometric model and numerical calculations of mean-first passage time

To gain a better understanding of the geometric protein-DNA binding model, we numerically compute the expected dissociation rates for a corresponding theoretical model using standard statistical mechanical methods. Within the stochastic kinetic model described by Fig. 1A and the equations above, we calculate the average protein dissociation rate as the inverse of the mean first-passage time (22, 91):

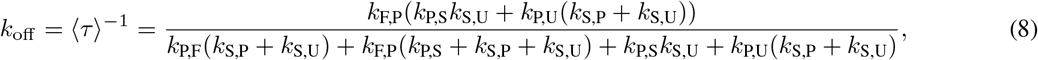

where *k*_A,B_ is the transition rate from state A to state B. As above, *k*_F,P_ and *k*_P,S_ depend on force. In addition, *k*_P,S_ is proportional to the concentration of competitor protein. The other rates are constant with the same values as in the simulation model.

The expected dissociation rate is given by Eq. 8. To use that expression, we must calculate the mean rates of partial dissociation, 〈*k*_F,P_〉, and competitor-binding-mediated saturation, 〈*k*_P,S_〉. These are given by averaging with respect to the equilibrium distribution of local DNA bending angles, which is weighted by the potentials of the geometric model above and the force that tends to align the DNA polymer along the tension axis. Thus, the partial dissociation rate is given by:

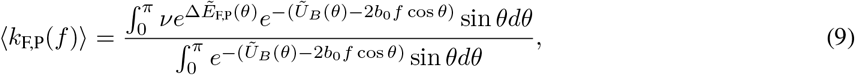

where *ν* = 1/*τ*_MC_ is the attempt rate for partial unbinding, which is chosen to match the simulations. Similarly, the competitor binding (saturation) rate is given by:

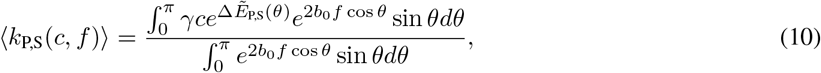

where *c* is the concentration of competitor proteins and *γ* is the binding attempt rate, which is again chosen to match the simulations.

We numerically compute these average rates and insert them into Eq. 8 to obtain the mean dissociation rate. All rate constants are reported as non-dimensionalized quantities. Rate constants are normalized by the rate constant with the same parameters (*x*_F,P_, *x*_P,S_, and *θ*_0_) calculated for *f* = 0 and *c* = 100 *μ*M. This rescaling mimics the adjustable energy barriers in Eqs. 6 and 7 in a simplified manner. As in simulations, we implement a cutoff to identify force-insensitive kinetic regimes; for the theory, *σ* = 0.001. Theoretical results are shown for *θ*_0,F_ = *θ*_0,S_ = *π*/3 and *x*_P,S_ = 1, unless noted.

## Results

### Geometric model coupling force and dissociation

The four-state model described above allows proteins to unbind via two different dissociation pathways, which are illustrated in Fig. 1B. The first is a spontaneous dissociation pathway, in which a protein (black dimer) sequentially transitions from fully bound to partially bound to unbound spontaneously, without the participation of competitor protein (gray dimer). The second dissociation pathway is the concentration-dependent facilitated pathway, where a protein transitions from fully bound to partially bound, and then a competitor protein binds to the partially vacated binding site and assists the original protein in unbinding. Fully bound and saturated proteins physically interact with the DNA substrate by bending the DNA polymer. This changes the local equilibrium bond angle, *θ*_0,A_. There are two force dependencies included in the models, which are governed by the force sensitivity parameters, *x*_F,P_ and *x*_P,S_ (see methods section above and Fig. 1A). In general, these two force dependencies enable the applied force, *f*, to accelerate the F-P transition and inhibit the P-S transition. By varying the force dependencies of the rates associated with the two different pathways, we observe a range of bond dissociation behaviors, described below.

### Dissociation rates in limiting scenarios

To demonstrate the main phenomenological features of our model, we first investigate the simplest scenarios, in which the system has only one force dependence (*i.e*., one of either *x*_F,P_ > 0 or *x*_P,S_ > 0). In these cases, the net protein dissociation rate depends in a simple manner on the single force-dependent transition rate (either *k*_F,P_ or *k*_P,S_).

#### Force-dependent spontaneous dissociation

We first consider the protein-DNA binding model with the force dependence restricted to only the spontaneous dissociation pathway. Thus, we vary *x*_F,P_ while fixing *x*_P,S_ = 0 so that only the unbinding transition from fully bound to partially bound depends on force. Here, the model demonstrates canonical slip bond behavior; the bond is weakened as more force is applied, so the bond lifetime decreases. This is manifested in the model by an observed increase in the dissociation rate, 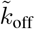. The force dependence of partial unbinding models the force dependence of observed “microdissociation” events, which is observed, for example, in experiments with HMGB and NHP6A (90).

As force is applied to the DNA polymer, it straightens, which decreases the local bending angle, *θ*. This frustrates the fully bound protein conformation, which has binding energy proportional to (*θ* – *θ*_0,F_), with *θ*_0,F_ > 0 (Eq. 7). Any *x*_F,P_ > 0 for both the theoretical and simulation model incorporates this decrease in the bending angle in the F to P unbinding step, leading to a more favorable unbinding step at high forces. This leads faster protein dissociation as force is increased, and results in slip-bond behavior in the models. This is true even when competitor proteins facilitate dissociation (in a force-independent manner), as shown in Figs. 2A and S1A.

**Figure 2:**
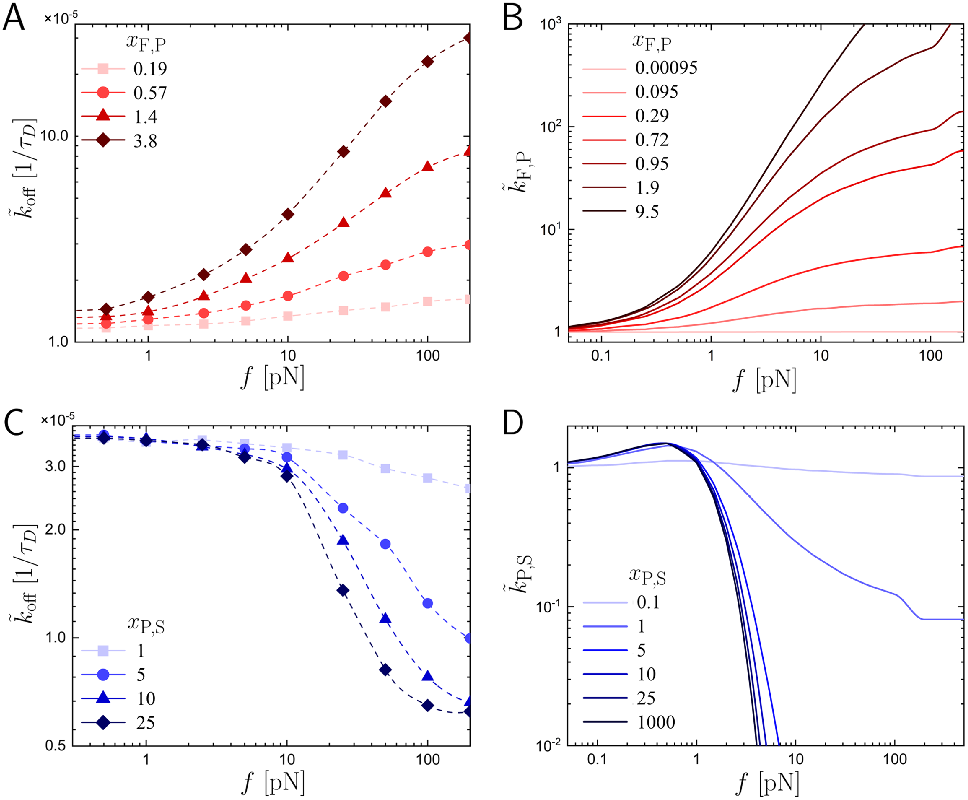
**A**: In simulations with force dependence only for partial dissociation (*i.e*., *x*_F,P_ > 0 and *x*_P,S_ = 0), the overall dissociation rate, 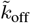, increases with force, *f*. The force dependence strengthens with increasing force sensitivity, *x*_F,P_ (light to dark red). **B**: The normalized partial dissociation rate, 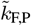, (which is different from the overall 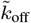 shown in panel A), calculated from Eq. 9. As *x*_F,P_ increases, the partial dissociation rate increases; this underlies the slip-bond behavior observed in simulations. **C**: In simulations with force sensitivity only for competitor binding (*x*_F,P_ = 0 and *x*_P,S_ > 0), the dissociation rate, 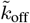, decreases with increasing force. As the force sensitivity, *x*_P,S_, increases (light to dark blue), the force dependence of the dissociation rate becomes sharper and the onset of strong force dependence shifts to lower forces. **D**: The normalized competitor binding rate, *x*_P,S_ (different than *k*_off_ shown in panel C) calculated from Eq. 10. For large enough forces, 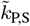 is generally suppressed by increasing force, which results in catch-bond kinetics. As *x*_P,S_ increases, *k*_P,S_ is suppressed more sharply and for smaller *f*. Results in all panels are for *c* = 100 *μ*M. Simulation results (A and C) are for *θ*_0,F_ = *θ*_0,S_ = *π*/6 and theoretical results (B and D) are for *θ*_0,F_ = *θ*_0,S_ = *π*/3. Note that the larger equilibrium angle in the theoretical model reduces excluded volume effects (also see Fig. S2), while maintaining similar kinetics to the simulations with a smaller equilibrium angle.

As *x*_F,P_ increases, the total dissociation rate becomes more force sensitive. As illustrated by the transition rate numerically calculated from Eq. 9, shown in Fig. 2B, the spontaneous dissociation rate, 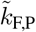 increases more rapidly with force for larger *x*_F,P_. Thus, in scenarios in which only partial dissociation depends on force, tension within DNA may accelerate dissociation of multivalent DNA-bending proteins.

#### Force-dependent facilitated dissociation

We also investigate the case in which only the facilitated dissociation pathway depends on force, *x*_P,S_ > 0 and *x*_F,P_ = 0; this leads to catch-bond behavior. This force dependence increases the energy barrier for a competitor protein to bind to a partially occupied DNA site. This models the inhibition of binding by DNA-bending proteins when DNA bending is inhibited by tensile forces (66, 87).

Applied forces in this scenario decrease the total dissociation rate by suppressing competitor binding, and thus slowing protein dissociation via the facilitated pathway. In the model, force decreases the average bending angle 〈*θ*〉. This increases the energy barrier for competitor protein binding, which is proportional to (*θ* – *θ*_0,S_)^2^. As shown in Fig. 2C, for finite competitor concentrations, the off rate, 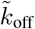, decreases with increasing force, *f*, in simulations, as expected for a catch bond.

For sufficiently high forces, *f*, and force sensitivities, *x*_P,S_, the off rate is insensitive to force, as seen for *f* > 100 pN and *x*_P,S_ = 25. For even sharper force dependencies (larger *x*_P,S_), competitor binding should be suppressed for any *f* > 0. Under such conditions, the facilitated dissociated pathway is so strongly inhibited by force that proteins can effectively only dissociate through the spontaneous pathway, leading to a force-independent *k*_off_. This trend can be inferred from the observed shift with increasing *x*_P,S_ in simulations (from light to dark blue) and for large *x*_P,S_ in the theory (Fig. S1B). The numerically calculated transition rate, *k*_P,S_, shown in Fig. 2D illustrates the changes in the facilitated rate with *x*_P,S_ and *f* that govern the total dissociation rate. The theoretical *k*_P,S_ is mostly consistent with simulations, but contrary to expectations from the simulations, the calculated rate is non-monotonic in force, increasing for small *f*. This occurs in the theoretical model because low forces shift the angular distribution toward the equilibrium angle (for *θ*_0,S_ < *π*/2), which enhances the competitor binding rate. In simulations, this effect is suppressed by excluded volume interactions, which reduce the mean bending angle at zero force. However, excluded volume can be incorporated into a more detailed version of the theoretical model (Fig. S2).

The two distinct variable force dependencies in the model thus have opposing effects on the total dissociation rate. This kinetic behavior is generic; for example, it can be reproduced by a simple theoretical model with purely exponential force dependencies (see Supplemental Text). However, in contrast to these limiting scenarios, physiological protein dissociation may have multiple force-dependent kinetic barriers. These force dependencies have differing degrees of control over the protein dissociation kinetics in different physical regimes, described below.

### Multivalent protein dissociation with multiple force dependencies

While the effect of force on each dissociation pathways is straightforward when considered independently, we observe more complex behavior in scenarios in which both pathways are force dependent. In this case, inhibition of competitor protein binding by force suppresses the facilitated pathway, which competes with the force-enhanced partial unbinding that accelerates the spontaneous dissociation pathway.

As we observe in Fig. 3A and B, this competition between the two pathways leads to three classes of bonds: the slip and catch bonds discussed above, and a hybrid bond we refer to as a “delayed catch bond.” The delayed catch bond is characterized by an off rate with a slip-bond force dependence for low forces and a transition to catch-bond behavior with increasing force; hence the onset of the catch bond is delayed as a function of force. At very high forces, the off rate may either increase with force (reverting to a slip bond) or be essentially insensitive to force. Fig. 3A and B show the transitions between catch, delayed catch, and slip bonds as a function of the force sensitivity ratio, *x*_P,S_/*x*_F,P_ for the particular parameter choice *x*_P,S_ = 10 for the simulations and *x*_P,S_ = 1 for the theory; other choices of *x*_P,S_ lead to qualitatively similar results (*e.g*., see Fig. S3A-B).

**Figure 3:**
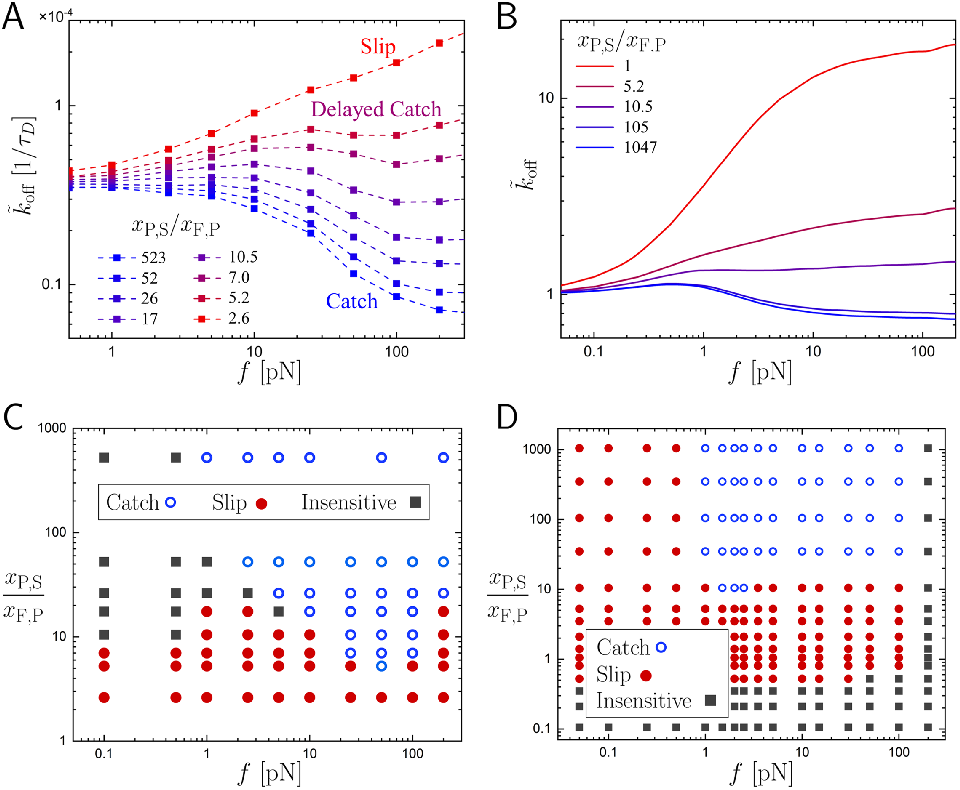
**A:** Simulation off rates, 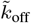, as a function of force, *f*, and sensitivity ratio, *x*_P,S_/*x*_F,P_, at *x*_P,S_ = 10 and *c* = 100 *μ*M. As the sensitivity ratio decreases (by increasing *x*_F,P_), dissociation kinetics change from catch-bond (blue) to delayed-catch-bond (purple and dark red) to slip-bond (red) behavior. **B:** Theoretical off rates as a function of force, *f*, and sensitivity ratio, *x*_P,S_/*x*_F,P_, at *x*_P,S_ = 1 and *c* = 100 *μ*M. Similar to the simulation data, as the ratio *x*_P,S_/*x*_F,P_ decreases, the force-dependent dissociation of the protein-DNA complex crosses over from catch-bond to slip-bond kinetics. **C:** A phase diagram in the ratio-force plane, constructed from the simulation data, shows physical regimes of decreasing *k*_off_, corresponding to catch bonds (open blue circles), and increasing *k*_off_, corresponding to slip bonds (solid red circles). **D:** The theoretical diagram shows regimes with catch-bond, slip-bond, and force-insensitive-bond kinetics (open blue circles, solid red circles, and solid gray squares, respectively). The theory agrees qualitatively with simulations, and reveals an additional force-insensitive regime for very low sensitivity ratios, *x*_P,S_/*x*_F,P_.

For low sensitivity ratios, the force dependence of the facilitated dissociation pathway is weak compared to that of the spontaneous pathway, whereas for high sensitivity ratios, the force dependence of the facilitated pathway is stronger. In both simulations and theory, for low sensitivity ratios, the protein-DNA complex acts as a slip bond, dissociating more rapidly with increasing force (red lines). As before, this occurs because force stimulates spontaneous dissociation while having relatively little effect on the competitor protein binding required for facilitated protein unbinding (*e.g*., in Fig. 2, compare slopes of dark lines in B to light lines in D).

As the sensitivity ratio is increased, the force has a greater inhibitory effect on the facilitated dissociation pathway. For moderate sensitivity ratios, this manifests itself at ~ 1 – 10 pN forces in the model as a change from an increasing to a decreasing off rate (dark red and purple lines in Fig. 3A and B). The onset of catch-bond kinetics is delayed in this scenario because the spontaneous pathway has a non-negligible force dependence, while the competitor binding rate, 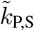, in the facilitated pathway is less sensitive to small forces. This disparity arises because *k*_F,P_ depends purely exponentially on the bending angle, *θ*_0,F_, whereas the competitor binding rate has a broad (*x*_P,S_-dependent) peak around the preferred binding angle, *θ*_0,S_ (see Eqs. 6 and 7). Moreover, at low forces, by accelerating the transition from fully to partially bound, the facilitated dissociation pathway is also effectively accelerated because the protein-DNA complex is more frequently susceptible to invasion by a competitor protein.

For very high sensitivity ratios, the force dependence of the facilitated pathway is dominant. Here, we observe catch-bond kinetics in the simulation, while the theoretical model predicts that the onset of the catch bond is delayed to higher forces (blue lines in Fig. 3A and B). This discrepancy occurs because the Gaussian chain approximation used in the theoretical model results in a small enhancement in the rate 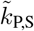 for small forces (Fig. 2D). This effect is suppressed in simulations by the additional chain stiffness imparted by excluded volume interactions. Nonetheless, we observe predominantly catch-bond kinetics in this regime because the facilitated dissociation pathway is far more sensitive than the spontaneous pathway to the applied force.

To characterize the broader force-dependent dissociation kinetics of this model, we measure the derivative of the dissociation rate for a range of sensitivity ratios, *x*_P,S_/*x*_F,P_, and forces, *f*. Fig. 3C and D shows the resulting phase diagrams for the simulations and theory. Here, we observe three types of force-dependent dissociation kinetics: force-insensitive bonds (gray squares), slip bonds (solid red circles), catch bonds (open blue circles). Combinations of these types of kinetics result in the three general classes of bonds described above.

For high force sensitivity ratios, where *x*_P,S_ ≫ *x*_F,P_, we always observe catch-bond kinetics in simulations (*i.e*., decreasing 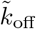). In contrast, the theoretical model predicts that the onset of catch-bond kinetics should arise at nonzero force for all sensitivity ratios. Again, this difference is due to the Gaussian chain approximation utilized in the theoretical model.

For lower force sensitivity ratios, we observe a growing domain of slip-bond behavior as *x*_P,S_/*x*_F,P_ decreases, and the transition to catch-bond kinetics occurs at higher forces. This occurs because the force dependence of competitor binding is weaker than that of partial dissociation. Below a critical sensitivity ratio, the force dependence of partial protein dissociation completely dominates, and only classical slip-bond kinetics are observed. For very low sensitivity ratios, below the minimum value tested in simulations, the theory predicts force-insensitive kinetics (Fig. 3D). In this regime, *x*_F,P_ is large and the spontaneous pathway is so sensitive to force so that partial dissociation occurs almost instantly for any finite *f*. Thus, even though the facilitated pathway remains sensitive to force (via *x*_P,S_), it no longer significantly accelerates dissociation; this is because because the rebinding transition (P to F) that is prevented by competitor binding no longer delays protein dissociation.

For all sensitivity ratios, the theoretical model predicts a force-insensitive domain for sufficiently high forces. This occurs because the facilitated pathway is completely suppressed by force and the spontaneous pathway cannot be further enhanced by additional application of force. In the simulations, this predicted force-independent regime likely occurs at forces above the maximum simulated *f*.

In summary, in our protein dissociation model, several varieties of force-dependent kinetics are possible. This behavior is largely controlled by the force sensitivity ratio, which determines the relative strength of the force dependence of the facilitated dissociation pathway as compared to that of the spontaneous dissociation pathway. The competition between the force dependencies of these two dissociation pathways leads to hybrid kinetic behaviors (“delayed catch bonds”), or in unbalanced cases, distinct limiting scenarios (slip or catch bonds).

### Effects of binding geometry

A key determinant of protein-DNA interactions is the geometry of the binding interface. Critically, this local geometry can determine the strength and nature of the coupling between applied forces and dissociation kinetics. To study how geometric factors may alter the force dependence of proteins undergoing facilitated dissociation, we varied the parameter *θ*_0,A_, which sets the equilibrium DNA bending angle for protein binding (Eq. 3). Alterations to *θ*_0,A_ thus regulate how the applied force, *f*, changes the overall dissociation rate, 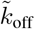.

We consider three different values of *θ*_0,F_ = *θ*_0,S_ ≡ *θ*_0_ in the simulation model: *π*/6, *π*/3, and *π*/2. We measure the dissociation rate, 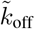, at different forces, *f*, and force sensitivity ratios, *x*_P,S_/*x*_F,P_ (fixing *x*_P,S_ = 10 and *θ*_0,U_ = *θ*_0,P_ = 0 for simplicity). Contour plots showing the dissociation rates for *θ*_0_ = *π*/6, *π*/3, and *π*/2 are shown in Fig. 4A-C, respectively. For all three cases, we see the same qualitative dissociation kinetics discussed above: catch bonds when the facilitated pathway is most sensitive to force (large *x*_P,S_/*x*_F,P_), slip bonds when the spontaneous pathway is most sensitive to force (smaller *x*_P,S_/*x*_F,P_), and delayed catch bonds when the force dependencies of the two pathways compete (intermediate *x*_P,S_/*x*_F,P_). These qualitative observations also hold for the theoretical model (Fig. S4).

**Figure 4:**
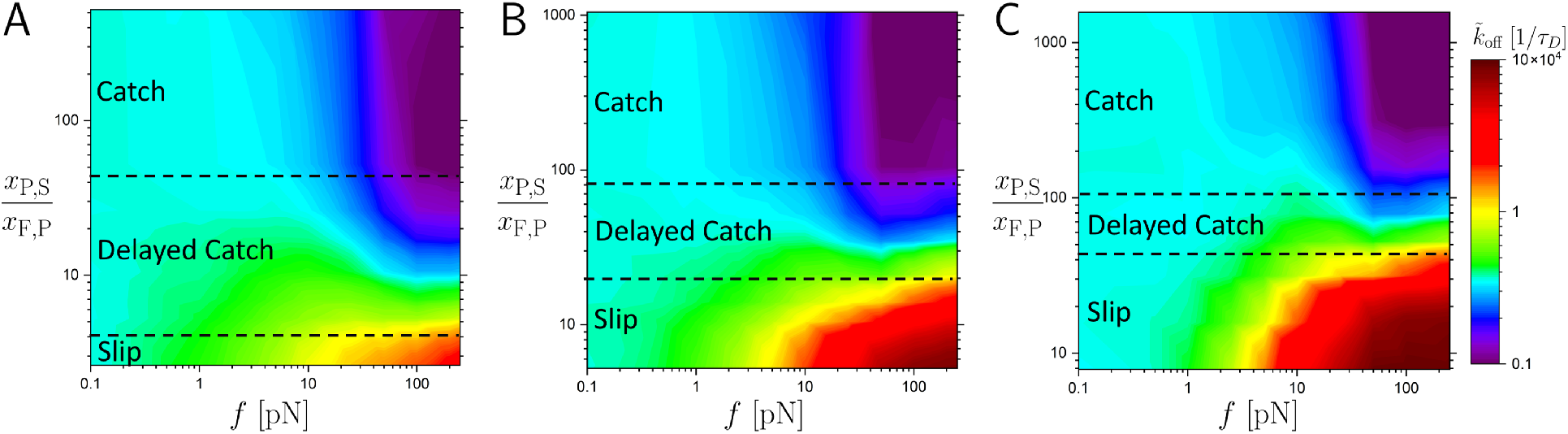
Contour plots of 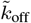 as a function of applied force, *f*, and sensitivity ratio, *x*_P,S_/*x*_F,P_, forbending angles *θ*_0,F_ = *θ*_0,S_ ≡ *θ*_0_ = *π*/6, *π*/3, and *π*/2 (**A-C**, respectively) with *x*_P,S_ = 10 and *c* = 100 *μ*M. Approximate boundaries between catch bond, delayed catch bond, and slip bond regimes are denoted by black dotted lines. Large off rates, corresponding to faster dissociation, are red, while small off rates (slower dissociation) are purple.

Nonetheless, binding geometry has a marked impact on the transitions between these dissociation behaviors. Most notably, as *θ*_0_ is increased, the range of force sensitivity ratios, *x*_P,S_/*x*_F,P_, in which delayed catch bonds are observed shrinks. This effect occurs because protein-DNA complexes with a large equilibrium DNA-bending angle are most strongly impacted by the applied force; straight DNA conformations induced by applied force have an energetic cost for protein binding that increases with *θ*_0_ (see Eqs. 6 and 7). Consequently, for large preferred bending angles, even small forces can induce sharp changes to the partial dissociation and competitor binding rates. Thus, the regime in which the two pathways compete narrows for highly bent interfaces because their respective force-dependent rates, *k*_F,P_ and *k*_P,S_, vary over different narrow force scales.

In addition to qualitative changes in the transitions between different regimes, we observe quantitative differences in the off rate as a function of *θ*_0_. In Fig. 4, purple regions of the contour plots indicate a slow 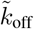, while red regions indicate fast 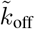. For larger bending angles, the red region extends to lower forces, which indicates that slip dissociation can be rapid even for small forces because of the mechanical energetic cost for the protein to remain bound to DNA. In contrast, 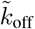 for the catch-bond regime is not drastically altered because it is minimized as the facilitated pathway is suppressed by force.

Altogether, altering the binding geometry by varying *θ*_0_ changes the degree of coupling between mechanics and kinetics. This coupling modulates the energy barriers between different binding states by determining the energy required for the protein to bend DNA. Thus, the binding geometry is a critical factor in regulating the competition between dissociation pathways and determining the force dependence of the overall protein dissociation rate.

### Effects of competitor concentration

Since the phenomenon of facilitated protein dissociation is characterized by concentration-dependent off rates, we study the force-dependent kinetics at different ambient concentrations of competitor molecules. At zero force, as the competitor concentration, *c*, increases, more initially bound proteins dissociate via the facilitated dissociation pathway. We thus hypothesized that increasing *c* could lead to the facilitated pathway becoming dominant for small forces and force sensitivity ratios; in turn, catch-bond kinetics would become more prevalent.

We consider the two limiting scenarios of high and low sensitivity ratios, which respectively, exhibit catch- and slip-bond kinetics at *c* = 100 *μ*M. We investigate how changing the concentration from ≈ 1 *μ*M to > 10^4^ *μ*M modulates dissociation behavior. Plots showing 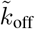 as a function of force, *f*, and concentration, *c*, are shown in Fig. 5A-D for simulations (A and B) and theory (C and D). Their corresponding bond phase diagrams are shown in Fig. 5E-H.

**Figure 5:**
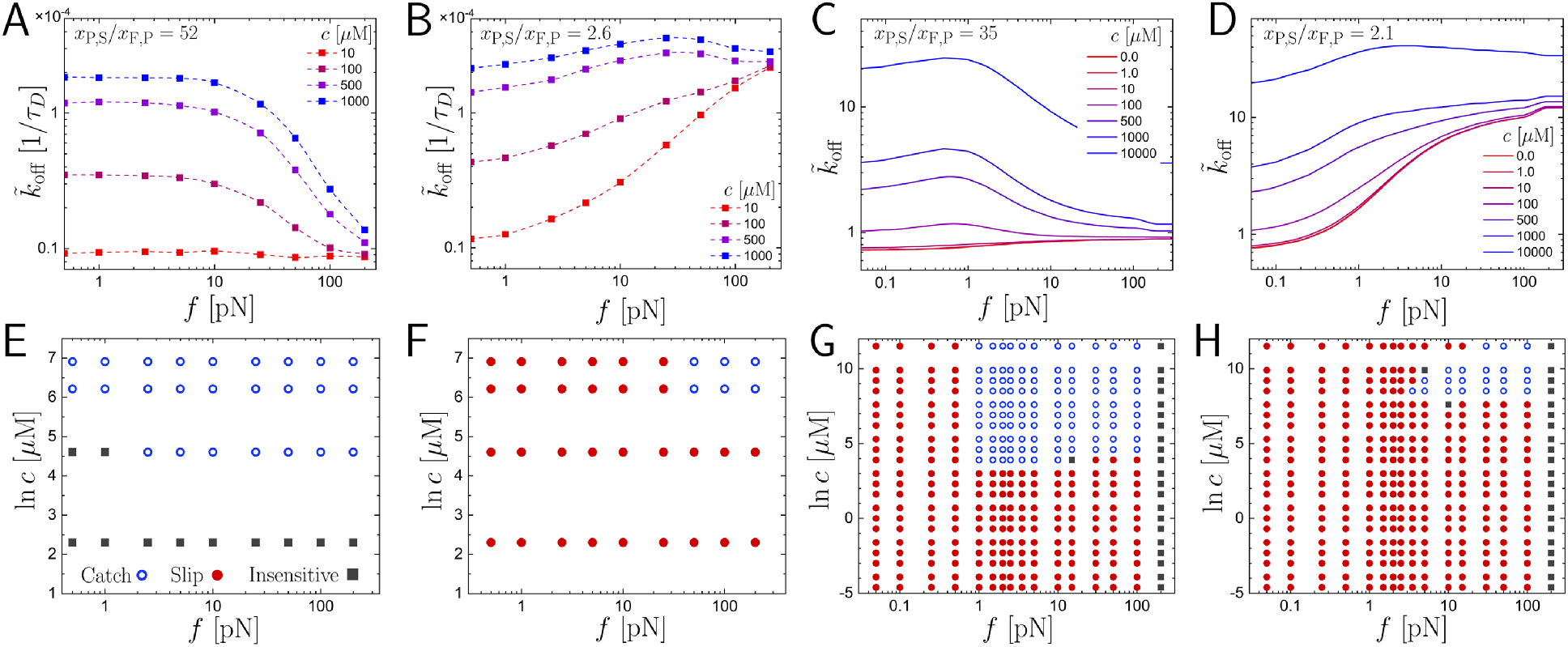
**A-B:** Simulation off rates, 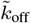, as a function of force, *f*, at various concentrations, *c*, where *θ*_0,F_ = *θ*_0,S_ = *π*/6. The simulation results shown are for sensitivity ratios of *x*_P,S_/*x*_F,P_ = 52 and 2.6, respectively. Different colored curves show different competitor concentrations, ranging from low (red) to high (blue). **C-D**: 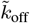 for the theoretical models, where *θ*_0,F_ = *θ*_0,S_ = *π*/3. The theoretical sensitivity ratios are *x*_P,S_/*x*_F,P_ = 35 and 2.1, respectively. **E-H:** Corresponding phase diagrams indicating the bond type as determined by *∂k*_off_/*∂f*, plotted in the concentration-force plane. Each panel correspond to the 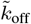 plot directly above it.

We first study the effect of varying competitor concentration on dissociation with a large force sensitivity ratio. For concentrations spanning at least three orders of magnitude in simulations, we observe only catch bonds or, at low *c*, force-insensitive bonds, as shown in Fig. 5A and E. The spontaneous pathway depends on force so weakly that almost completely irrespective of concentration, the facilitated pathway is more sensitive to force. For high concentrations (blue lines), this leads to a 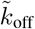 that decreases with force, *i.e*., a catch bond. For lower concentrations, *c* ≤ 10 *μ*M (red lines), there are very few competitor proteins available to participate in facilitated dissociation. So, while the facilitated dissociation pathway is sensitive to force, DNA-bound proteins rarely dissociate via this pathway, which leads to a nearly force-independent off rate.

As with varying the force sensitivity ratio, these observations hold for the theoretical model with the caveat that we observe delayed onset catch bonds instead of catch bonds (Fig. 5C and G). Again, this is due to discrepancies between the low-force DNA conformations, and thus, competitor binding rates, found in the different models. Fig. S3C and D display the same trends for the theoretical model with a different force sensitivity for competitor binding, *x*_P,S_ = 10. The theoretical model also allows us to observe the slip-bond regime that is present for very low competitor concentrations. Because competitor proteins are scarce in this regime, the effect of the facilitated dissociation pathway is negligible, and its force dependence is unimportant.

We also investigate the alternative limiting case in which the force sensitivity ratio is small, so that the spontaneous pathway is more sensitive to force. Here, for low concentrations (*c* = 10 *μ*M in simulations, red lines in Fig. 5B), we find pure slip-bond kinetics due to the force dependence of the partial unbinding transition. This is expected because there are very few proteins to facilitate the dissociation of the initially bound proteins, so variations in competitor binding rates contribute negligibly to the total dissociation rate. Moreover, because *x*_F,P_ is larger in this scenario, the slip regime can be observed in the simulations for small *c* (Fig. 5B and F). For sufficiently high concentrations (*c* ≥ 500 *μ*M), we see delayed catch bonds. With these high competitor concentrations (blue and purple lines in Fig. 5B), proteins can easily dissociate via the facilitated pathway due to the frequency with which competitor proteins attempt to invade the DNA binding site. While force-induced inhibition of competitor binding is outweighed by force-enhanced partial dissociation pathway at low forces, for large forces, competitor proteins are strongly inhibited from binding. This shuts off a major dissociation pathway, which leads to the emergence of catch-bond behavior.

The theoretical calculations with low sensitivity ratio, shown in Fig. 5D and H and Fig. S3E and F, also reveal a change from slip-bond dissociation at low concentrations to delayed-catch-bond kinetics at high concentrations. Additionally, in the theoretical model, we observe the high-force convergence of the off rates at different concentrations (Fig. 5D), as in the simulation data (Fig. 5B). This convergence occurs because the facilitated dissociation pathway is completely suppressed by large forces, and thus concentration does not regulate the off rate in the high-force regime.

These results indicate the fundamental role that competitor protein (or biomolecule) concentration plays in regulating force-dependent unbinding kinetics. Concentration sets the overall magnitude of the facilitated dissociation effect and is thus an essential ingredient of the catch bond exhibited by the model.

## Discussion

DNA-binding proteins may exhibit catch-bond unbinding kinetics when subjected to force, provided that two conditions are met: 1) the protein can undergo facilitated dissociation and 2) protein binding is inhibited by applied tension. The first condition can be met by proteins that form multiple contacts with DNA, so that they may partially unbind and allow a competitor protein to invade the DNA binding site and thus, accelerate unbinding. The second condition will generally be met when a protein binds DNA in a configuration that is suppressed by tension, such as in a bent or overtwisted conformation.

Here, we investigated the effects of several key physical variables on force-dependent facilitated dissociation of proteins from DNA using both simulation and theory. With two force-dependent pathways for protein dissociation – spontaneous and facilitated – we observe three qualitatively distinct types of unbinding kinetics. Generically, catch bonds are observed when the facilitated pathway is strongly inhibited by applied force. Force suppresses competitor binding to DNA, which impedes the facilitated dissociation pathway (Fig. 2C and D). Slip-bond behavior, in contrast, is observed in cases in which the spontaneous pathway is more sensitive to force, so the complex exhibits canonical force-dependent dissociation kinetics (*e.g*., Fig. 2A and B). Between these limiting scenarios, the force dependencies of the two pathways compete with each other. Here, we may observe delayed catch bonds, in which 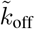 as a function of *f* initially increases, then decreases, and finally increases once again (Fig. 3A and B). More generally, the observable types of force-dependent dissociation kinetics depend on quantitative details of the system of interest. In particular, other transition rates in the process shown in Fig. 1 could depend on force, which could expand the range of dissociation behaviors considerably. Beyond the force sensitivities, we have found that several other factors – both intrinsic and extrinsic – regulate force-dependent dissociation kinetics of multivalent DNA-binding proteins (Figs. 4 and 5). Below, we discuss these factors, experimental evidence and tests for force-dependent facilitated dissociation, and the possible physiological relevance of our results.

### Multivalent proteins can form catch bonds via extrinsic factors

We have found the surprising behavior that protein-DNA complexes dissociating due to competition with other proteins in solution may dissociate more slowly when subjected to force. Thus, these complexes form “catch bonds,” with force-dependent kinetics qualitatively similar to that of other proteins and complexes, including cell adhesion proteins and molecular motors (69, 70, 76, 78, 79, 81–84). However, the catch bond we observe in this protein-DNA system differs from other complexes that are known to exhibit catch-bond behavior. Many catch bonds occur due to intrinsic factors, such as conformational changes within the protein-ligand complex, as is the case with the bacterial adhesion protein FimH (75, 79). Catch bonds can also arise when highly elastic transition states occur in *e.g*. protein unfolding, due to the conformational entropy of unfolded protein strands (92–94). With facilitated dissociation, the observed catch bond is due to force-inhibited binding of a third competitor molecule, while the original protein-DNA complex in the absence of these competitor molecules actually demonstrates the classical slip-bond kinetics. Thus, whereas the change in force-induced affinity in FimH is *intrinsic* to the molecular structure of the bond, the DNA-binding proteins that we consider undergo *extrinsic*, force-induced changes in affinity due to the binding kinetics of competitor molecules.

The extrinsic mechanism in this work is conceptually similar to a two-pathway models that have been proposed to describe other proteins that exhibit intrinsic catch-bond dissociation kinetics (13, 69–73). In those models, at low forces, a fast dissociation pathway dominates the energy landscape. However, as force is increased, the energy barrier for proceeding through a second, slower pathway is decreased. This results in longer total dissociation rates and catch-bond kinetics. The “fast” and “slow” pathways are analogous to the facilitated and spontaneous dissociation pathways, respectively. Different intrinsic factors responsible for driving the enhancement of the slower pathway have been proposed, including force-dependent, molecular-level interactions and multiple intermediate conformational states that differ in thermodynamic stability based on the force regime (13, 69, 71, 73). The extrinsic mechanism explored in this work is regulated by a broader collection of physical variables.

Among these variables, the competitor protein concentration, an extrinsic factor, is essential to drive facilitated dissociation; thus, it necessarily determines the nature of the overall force-dependent kinetics that we observe (Fig. 5). Low concentrations generally lead to only slip-bond kinetics due to the limited role of the facilitated dissociation pathway. As concentration is increased, if both the spontaneous and facilitated pathways are sensitive to force, we observe a transition from a slip bond to a delayed catch bond because the facilitated and spontaneous dissociation pathways compete. When the spontaneous pathway has a very weak force dependence or the facilitated pathway has a very strong force dependence, we may observe only catch bonds.

### The shape of the binding interface couples to force-dependent dissociation kinetics

The preferred binding geometry of the protein on DNA, and thus the local geometry of the binding site is an important intrinsic factor that governs protein dissociation. The binding geometry is governed by two types of parameter in our model, the equilibrium bending angles, *θ*_0,F_ and *θ*_0,S_, and the force sensitivities, *x*_F,P_ and *x*_P,S_ (Figs. 2 and 4).

The force sensitivities, which directly govern dissociation kinetics, implicitly describe the geometry of the protein-DNA complex. We interpret the force sensitivities as dictating the degree to which a bound protein deforms when the DNA substrate is bent away from the equilibrium binding angle. For large force sensitivities, the protein (and/or the DNA binding site) is stiff. With a high force sensitivity for the partial unbinding transition (governed by *x*_F,P_ in Eq. 7), the protein is not strongly distorted as DNA is straightened by applied force. As a result, the energy barrier for partial unbinding decreases, so the protein-DNA bond weakens and the corresponding transition rate, *k*_F,P_, increases. Similarly, the force sensitivity *x*_P,S_ that can inhibit competitor binding describes the degree to which a protein (and binding site) may deform in order to saturate a partially bound binding site. When this sensitivity is high, the protein is stiff and unlikely to fluctuate into a conformation that can bind a straightened segment of DNA. Thus, together, these force sensitivities implicitly describe conformational changes that are possible for the protein and corresponding DNA binding site.

The preferred angle for protein-DNA binding explicitly incorporates geometric factors into the model. In the model, the equilibrium bending angle directly controls how strongly the force sensitivity couples to the transition energy barriers, so geometry alters the overall force dependence in a manner similar to that of the sensitivity ratio. Larger equilibrium bends in the DNA (governed by *θ*_0,F_ and *θ*_0,S_ in Eq. 3) make both the facilitated and spontaneous pathways more sensitive to force (Eqs. 6 and 7). In turn, as both pathways become more sensitive to force, the crossover between the catch and slip bond kinetic regimes narrows, and each regime is expanded (Fig. 4). Because the force dependencies become so extreme, the system inevitably falls into one of the two limiting scenarios due to the imbalance in the force dependencies of the partial dissociation and competitor binding rates (*k*_F,P_ and *k*_P,S_, respectively).

For more complicated binding geometries, we expect correspondingly more complicated force-dependent dissociation rates. For instance, we included only two force-dependent steps out of the possible six in the four-state model. Including additional force dependencies, even in steps that are not associated with a geometric change, could significantly alter the dissociation kinetics. Each dissociation pathway (facilitated and spontaneous) would then have multiple cooperating or competing force-dependent steps, which could lead to a more diverse set of dissociation behaviors. Moreover, the functional forms of the force dependencies may govern the observed dissociation kinetics. Consequently, we anticipate that in biological manifestations of protein-DNA facilitated dissociation, the force-dependent kinetics will depend on structural properties of the protein and the local DNA/chromatin environment. Indeed, it has been observed that the dissociation rates of the transcriptional regulators CueR and ZntR in bacteria are sensitive to local chromatin conformation (30). Thus, while our simple four-state model illustrates the general phenomenology of force-dependent facilitated dissociation, forces may differentially govern kinetics through several biophysical mechanisms.

Intriguingly, these force-dependent kinetics may feedback into the local structure of the DNA substrate. Even at zero force, facilitated dissociation has been shown to regulate the conformation of stretched and twisted DNA (87). Thus, further studies of DNA interacting with proteins with force-dependent facilitated dissociation kinetics could lead to novel insights structure and dynamics of protein-bound DNA and chromatin.

### Single-molecule experiments to test the model

The force-dependent facilitated dissociation kinetics predicted in this work could be observed using established experimental setups. In particular, we propose that *in vitro* single-molecule experiments that mirror our model (Fig. 1B) would be able to measure the force dependence of facilitated dissociation. In the experiments, DNA is tethered to a surface at one end and to a bead at the other end (Fig. 1B). The bead is manipulated via optical or magnetic tweezers (*e.g*., (20, 28, 63, 65, 66, 88–90)), which can exert ~ 0.1 – 10 pN tensile forces on the tethered DNA.

To measure dissociation rates, protein initially bound to DNA should be fluorescently labeled, as in previous experiments studying facilitated dissociation of the bacterial nucleoid-associating protein Fis (20). Non-fluorescing competitor proteins may be flowed in so that as competitors from solution exchange with the initially bound fluorescent proteins, the tethered DNA strand becomes darker. Using an approach with distinguishable sets of protein is essential because simply measuring the number of bound proteins is insufficient to determine the dissociation rate because proteins may exchange, *i.e*., when a protein vacates a DNA binding site, the binding site may nonetheless remain occupied by the competitor protein (21–23, 30). However, with one species of “light” protein and one species of “dark” (competitor) protein, dissociation rates can be measured by observing the decay of the fluorescence signal along the tethered DNA. This experiment can probe kinetics at various tensile forces and competitor concentrations, and other physical variables, such as ionic concentration or composition, can be varied to tune transition energy barriers (22, 29, 65, 95) or preferred binding geometry (89).

A broad class of proteins could be tested in these experiments. Candidate proteins include those that both bend DNA and unbind from DNA by facilitated dissociation. A variety of proteins satisfy these criteria, and thus, in principle, could exhibit catch-bond kinetics via force-dependent facilitated dissociation. Specifically, we hypothesize that Fis or HU (20, 22, 28) could exhibit catch-bond kinetics while undergoing facilitated dissociation. Other possible examples include transcription factors such as CueR and ZntR, which undergo facilitated dissociation with rates that depend on the local chromatin conformation (30).

A variation of the above experiment with single-stranded DNA binding protein (SSB) suggests that facilitated dissociation can lead to catch-bond-like force-dependent kinetics (96). In this experiment, dissociation of SSB from its binding site is facilitated by “intersegment transfer,” *i.e*., DNA segments from the same DNA strand compete with binding by the protein to the initial binding site and eventually strip it from the binding site. In this experiment, force on the DNA suppresses the interaction of these distant DNA segments because DNA looping is inhibited by tension. Thus, force suppresses facilitated dissociation in this experiment, which is precisely what is required for the catch bond effect.

### Relevance to *in vivo* protein kinetics and function

Functionally, many catch bonds are complexes that must bear forces, such as adhesion proteins and molecular motors (69, 70, 76, 78, 79, 81–84). Indeed, some proteins that undergo facilitated dissociation are architectural proteins, such as the bacterial protein Fis, which can DNA stabilize loops against applied forces (63, 97). In addition, several recent studies suggest that facilitated dissociation may have important regulatory effects in living cells (23, 29, 30, 36, 98, 99). Our model suggests that these facilitated dissociation kinetics may be modulated by mechanical stresses transmitted through DNA/chromatin, and more generally, the local conformation of the genome.

Our main prediction is that forces can inhibit the dissociation of a DNA-bound protein by inhibiting competitor-mediated dissociation and exchange with biomolecules from the surrounding environment. It has previously been suggested that cells may utilize facilitated dissociation to efficiently regulate transcription through modulation of transcription factor binding and unbinding kinetics (23, 30, 98, 99). Our results suggest that large mechanical stresses within the genome would locally inhibit facilitated unbinding kinetics. Thus, transcriptional regulation by facilitated dissociation could be specifically targeted to mechanically relaxed regions of the genome.

It has also been shown that competitor DNA strands can facilitate protein dissociation by effectively stripping partially bound proteins from the protein-bound DNA (28, 63). For this process, competitor binding (P to S in Fig. 1A) should depend far less sensitively on the conformation of protein-bound DNA because competitor DNA binds the partially bound protein. Instead, competitor binding should depend on the conformation of and tension within the competitor DNA molecule. In contrast to the case of competitor-protein-mediated dissociation described above, this process should favor protein transfer to DNA segments with lower mechanical stress. This may be relevant, for instance, during transcription, in which RNA polymerase may exert ~ 10 pN forces (56, 57). There, it has been observed that nucleosomes initially proximal to the translocating polymerase are ejected, and subsequently found on nearby strands of DNA (100–104). This may be a prototypical example of force-dependent facilitated dissociation mediated by competitor DNA.

Force-dependent facilitated dissociation by competitor DNA may also regulate diffusive searches for target sites. For example, recent theoretical work suggests that protein target searches may be governed by the local chromatin conformation (105). Again, the force dependence of protein dissociation may either assist proteins searching for mechanically relaxed regions of chromatin or inhibit intersegment transfer as observed *in vitro* (96).

More generally, our work illustrates how mechanical forces may regulate facilitated protein-DNA dissociation. We anticipate that a wide variety of multivalent proteins may be subjected to these non-canonical, extrinsically regulated kinetics *in vivo*.

## Author contributions

KD, CES, and EJB designed the research and discussed all results. KD and CES developed the simulation model. KD and JZ performed and analyzed simulations. EJB developed the theoretical model and performed and analyzed the numerical calculations. CES supervised work by KD and JZ. KD and EJB wrote the manuscript with input from CES.

## Acknowledgements

KD, JZ, and CES acknowledge support from the National Institute Of General Medical Sciences of the National Institutes of Health under Award Number T32GM070421. The content is solely the responsibility of the authors and does not necessarily represent the official views of the National Institutes of Health. EJB was supported by the NSF Physics of Living Systems (15049420) and by the Center for 3D Structure and Physics of the Genome of the NIH 4DN Consortium (DK107980).

