## Supplementary text, tables, and figures for "Force-dependent facilitated dissociation can generate protein-DNA catch bonds"

July 8, 2019

#### Supplementary text

##### Simple theoretical model with different force dependencies

To illustrate the generality of the catch-bond effect generated by force-dependent facilitated dissociation, we consider a different stochastic kinetic model. This model assumes a purely exponential force dependence for each of the force-dependent transition rates, as in classical transition-rate theories (1, 2). We consider the reaction scheme given by Fig. 1A in the main text. The force dependent rates are

$$k_{\mathrm{F,P}} = \nu e^{-\Delta \tilde{E}_{\mathrm{F,P}} + f x_{\mathrm{F,P}}} \tag{1}$$

for partial unbinding (F $\rightarrow$ P) and

$$k_{\mathrm{P,S}} = \gamma c e^{-\Delta \tilde{E}_{\mathrm{P,S}} - f x_{\mathrm{P,S}}} \tag{2}$$

for competitor binding, *i.e.*, binding site saturation (P $\rightarrow$ S). Here, the force sensitivities, $x_{\mathrm{F,P}}$ and $x_{\mathrm{P,S}}$, have units of length per $k_B T$. Other transition rates are constant, as described in the main text, except that here, we omit the reverse reactions (P $\rightarrow$ F and S $\rightarrow$ P) in order to simplify the analysis.

Within this model, the net dissociation rate, $k_{\mathrm{off}}$, can have both slip- and catch-bond regimes, depending on the force sensitivities, $x_{\mathrm{F,P}}$ and $x_{\mathrm{P,S}}$. There are two particularly illustrative simple examples: 1) the case in which only partial dissociation depends on force ($x_{\mathrm{F,P}} > 0$ and $x_{\mathrm{P,S}} = 0$) and 2) the case in which only saturation depends on force ($x_{\mathrm{F,P}} = 0$ and $x_{\mathrm{P,S}} > 0$).

In case 1, the dissociation rate may be written as

$$k_{\mathrm{off}} = \frac{(k_{\mathrm{P,U}} + \gamma c)}{(e^{f x_{\mathrm{F,P}}} k_{\mathrm{F,P}} + k_{\mathrm{P,U}}) k_{\mathrm{S,U}} + (e^{f x_{\mathrm{F,P}}} k_{\mathrm{F,P}} + k_{\mathrm{S,U}}) \gamma c} k_{\mathrm{F,P}} k_{\mathrm{S,U}} e^{f x_{\mathrm{F,P}}}, \tag{3}$$

which manifestly increases with force, $f$. This occurs because partial dissociation is accelerated by force, while the remainder of the dissociation process is not affected. Thus, protein-DNA binding in this scenario acts as a slip bond, which dissociates more rapidly as tension increases.

In case 2, force suppresses the facilitated dissociation pathway by inhibiting competitor binding. The dissociation rate is

$$k_{\mathrm{off}} = \frac{k_{\mathrm{P,U}} + \gamma c e^{-f x_{\mathrm{P,S}}}}{(k_{\mathrm{F,P}} + k_{\mathrm{P,U}}) k_{\mathrm{S,U}} + (k_{\mathrm{F,P}} + k_{\mathrm{S,U}}) \gamma c e^{-f x_{\mathrm{P,S}}}} k_{\mathrm{F,P}} k_{\mathrm{S,U}}, \tag{4}$$

which manifestly decreases with increasing force, $f$, if $k_{\mathrm{S,U}} > k_{\mathrm{P,U}}$ (since $k_{\mathrm{off}}$ increases as the second term in the numerator, $\gamma c e^{f x_{\mathrm{P,S}}}$, increases, and we assume $x_{\mathrm{P,S}} > 0$). The overall dissociation rate decreases as force increases because it was previously dominated by the facilitated pathway, which is inhibited by the force.

Thus, the two protein dissociation pathways of proteins have qualitatively distinct force dependencies. This analysis illustrates that even simple force dependencies can lead to catch-bond kinetics for proteins undergoing facilitated dissociation. As in the models described in the main text, scenarios in which both $x_{\mathrm{F,P}} > 0$ and $x_{\mathrm{P,S}} > 0$ may result in different combinations of catch- and slip-bond kinetic regimes, depending on the strengths of the force sensitivities.


### Supplementary tables and figures

**Table S1:** Reference energy barrier values ($\Delta\tilde{E}_{\mathrm{b}}^{*}$) used in Eq. 6 in the main text. These values are found by parameterizing Eq. 6 to match all dissociation rates at $f = 0$. All other values in Eq. 6 are constant (other than the variable $\theta$, which is measured instantaneously in the simulation).

| $\theta_{0,\mathrm{S}}$ and $\theta_{0,\mathrm{F}}$ | $x_{\mathrm{P,S}}$ | $\Delta\tilde{E}_{\mathrm{b}}^{*}$ |
|---|---|---|
| $\pi/6$ | 1 | 1.2 |
| | 5 | 0.72 |
| | 10 | 0.4 |
| | 25 | -0.06 |
| $\pi/3$ | 1 | 0.1 |
| | 5 | -4 |
| | 10 | -9 |
| | 25 | -24 |
| $\pi/2$ | 1 | -2.6 |
| | 5 | -17.1 |
| | 10 | -35.5 |
| | 25 | -90 |

**Table S2:** Reference energy barrier values used in Eq. 7 in the main text. These values are found by parameterizing Eq. 7 to match all dissociation rates at $f = 0$.

| $\theta_{0,\mathrm{S}}$ and $\theta_{0,\mathrm{F}}$ | $x_{\mathrm{F,P}}$ | $\Delta\tilde{E}^*_{\mathrm{Ub}}$ |
|---|---|---|
| $\pi/6$ | $\leq 0.02$ | 10 |
| | 0.19 | 9.95 |
| | 0.38 | 9.88 |
| | 0.57 | 9.83 |
| | 0.95 | 9.84 |
| | 1.4 | 9.95 |
| | 1.9 | 10.16 |
| | 3.8 | 11.7 |
| $\pi/3$ | $\leq 0.25$ | 10 |
| | 0.29 | 10.01 |
| | 0.48 | 10.16 |
| | 0.72 | 10.5 |
| | 0.95 | 10.95 |
| | 1.9 | 14.35 |
| $\pi/2$ | $\leq 0.05$ | 10 |
| | 0.13 | 10.1 |
| | 0.19 | 10.17 |
| | 0.32 | 10.42 |
| | 0.64 | 11.54 |
| | 1.3 | 15.75 |

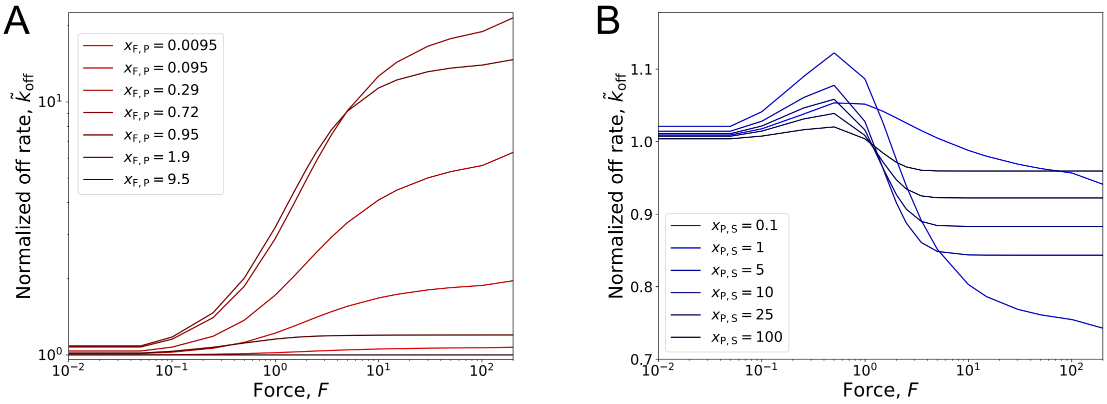


**Figure S1:** Normalized dissociation rates corresponding to theoretical models with a single force dependence (Fig. 2B and D in the main text). **A:** Normalized dissociation rate, $\tilde{k}_{\mathrm{off}}$, for the theoretical model with force-dependent partial dissociation ($x_{\mathrm{F,P}} > 0$, increasing from light to dark red) and force-independent facilitated dissociation ($x_{\mathrm{P,S}} = 0$). **B:** Normalized dissociation rate, $\tilde{k}_{\mathrm{off}}$, for the theoretical model with force-independent partial dissociation ($x_{\mathrm{F,P}} = 0$) and force-dependent facilitated dissociation ($x_{\mathrm{P,S}} = 0$, increasing from light to dark blue). Plots show results for $c = 100$ $\mu$M and $\theta_0 = \pi/3$.

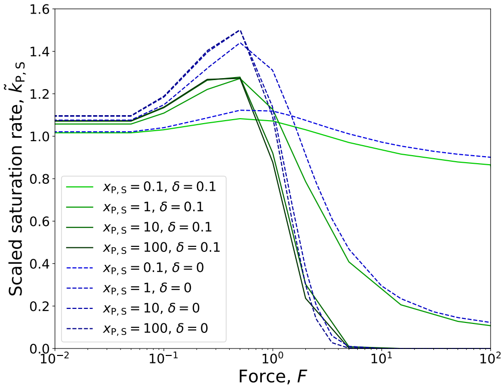


**Figure S2:** Scaled saturation rate, $\tilde{k}_{\mathrm{P,S}}$, for the theoretical model with an additional free energy term included to account for excluded volume effects. Here, the free energy is given by $\mathcal{F} = \tilde{U}_B(\theta) - 2b_0 f \cos\theta + \theta^2\delta$, where $\delta = 0.1$ governs the effective stiffness imparted by excluded volume. This expression is used to compute $\langle k_{\mathrm{P,S}}(c, f)\rangle$ as in Eq. 10 in the main text. Scaled rate with excluded volume is shown by the solid green lines, with increasing $x_{\mathrm{P,S}}$ from light to dark green. Results for the theory without excluded volume are shown for comparison by dashed blue lines (increasing $x_{\mathrm{P,S}}$ from light to dark blue). Bending stiffness imparted by excluded volume effects reduces the non-monotonicity of $\tilde{k}_{\mathrm{P,S}}$. Results are shown for $\theta_0 = \pi/3$.

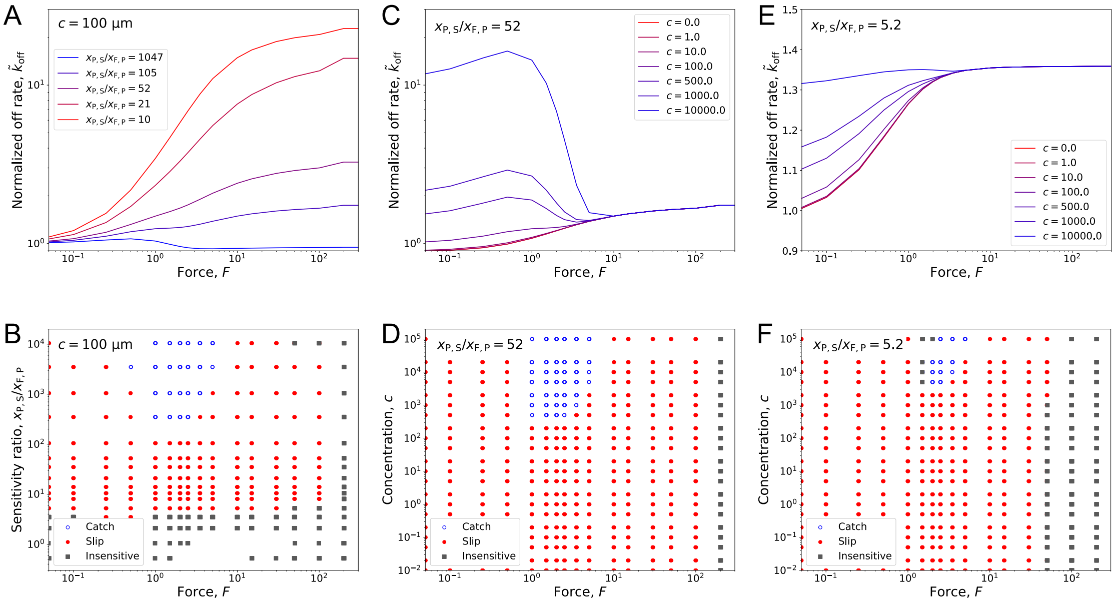


**Figure S3:** Theoretical results computed with the competitor binding force sensitivity fixed at $x_{\mathrm{P,S}} = 10$. **A:** Normalized off rate, $\tilde{k}_{\mathrm{off}}$, as a function of force, $f$, for different force sensitivity ratios, which are indicated by color (low ratio in red to high ratio in blue). Competitor concentration is fixed at $c = 100$ $\mu$M. **B:** Phase diagram for the kinetics corresponding to panel A. **C:** Normalized off rate, $\tilde{k}_{\mathrm{off}}$, as a function of force, $f$, for different concentrations, $c$, which are indicated by color (low concentration in red to high concentration in blue). Force sensitivity ratio is fixed to $x_{\mathrm{P,S}}/x_{\mathrm{F,P}} = 52$. **D:** Phase diagram for the kinetics corresponding to panel C. **E:** Normalized off rate, $\tilde{k}_{\mathrm{off}}$, as a function of force, $f$, for different concentrations, $c$, which are indicated by color (low concentration in red to high concentration in blue). Force sensitivity ratio is fixed to $x_{\mathrm{P,S}}/x_{\mathrm{F,P}} = 5.2$. **F:** Phase diagram for the kinetics corresponding to panel E. All panels show results for $\theta_0 = \pi/3$.

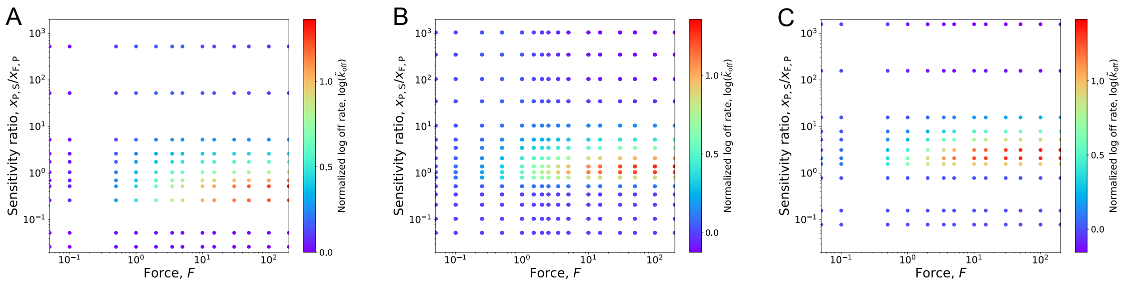


**Figure S4:** Normalized dissociation rates, $\tilde{k}_{\mathrm{off}}$ for various sensitivity ratios, $x_{\mathrm{P,S}}/x_{\mathrm{F,P}}$, and forces, $f$. Value of $\tilde{k}_{\mathrm{off}}$ is indicated by color from purple to red for slow to fast rates for **A:** $\theta_0 = \pi/6$, **B:** $\theta_0 = \pi/3$, and **C:** $\theta_0 = \pi/2$.
